# Recovery and analysis of transcriptome subsets from pooled single-cell RNA-seq libraries

**DOI:** 10.1101/408740

**Authors:** Kent A. Riemondy, Monica Ransom, Christopher Alderman, Austin E. Gillen, Rui Fu, Jessica Finlay-Schultz, Gregory Kirkpatrick, Jorge Paola Di, Peter Kabos, Carol A. Sartorius, Jay R. Hesselberth

## Abstract

Single-cell RNA sequencing (scRNA-seq) methods generate sparse gene expression profiles for thousands of single cells in a single experiment. The information in these profiles is sufficient to classify cell types by distinct expression patterns but the high complexity of scRNA-seq libraries often prevents full characterization of transcriptomes from individual cells. To extract more focused gene expression information from scRNA-seq libraries, we developed a strategy to physically recover the DNA molecules comprising transcriptome subsets, enabling deeper interrogation of the isolated molecules by another round of DNA sequencing. We applied the method in cell-centric and gene-centric modes to isolate cDNA fragments from scRNA-seq libraries. First, we resampled the transcriptomes of rare, single megakaryocytes from a complex mixture of lymphocytes and analyzed them in a second round of DNA sequencing, yielding up to 20-fold greater sequencing depth per cell and increasing the number of genes detected per cell from a median of 1,313 to 2,002. We similarly isolated mRNAs from targeted T cells to improve the reconstruction of their VDJ-rearranged immune receptor mRNAs. Second, we isolated *CD3D* mRNA fragments expressed across cells in a scRNA-seq library prepared from a clonal T cell line, increasing the number of cells with detected *CD3D* expression from 59.7% to 100%. Transcriptome resampling is a general approach to recover targeted gene expression information from single-cell RNA sequencing libraries that enhances the utility of these costly experiments, and may be applicable to the targeted recovery of molecules from other single-cell assays.

## INTRODUCTION

New methods that measure mRNA abundance in hundreds to thousands of single cells have been used to understand gene expression heterogeneity in tissues (Zheng et al. 2017; Macosko et al. 2015; Klein et al. 2015; Cao et al. 2017). But these single-cell RNA-seq experiments have a tradeoff: instead of surveying gene expression at great depth, they generate a sparse gene expression profile for each cell in a population. This information is often sufficient to identify cell types in a population, but provides only a glimpse of genes expressed in a given cell (Pollen et al. 2014). Moreover, mRNAs in each cell are captured stochastically, leading to false negatives in identification of expressed genes in many cells (Bacher and Kendziorski 2016).

Single-cell RNA-seq experiments can identify rare cell populations that have distinct gene expression profiles. Previous studies have identified retinal precursors (Shekhar et al. 2016; Macosko et al. 2015), hematopoietic stem cells (Olsson et al. 2016), rare immune cells (Yu et al. 2016), and novel lung cell types (Plasschaert et al. 2018) in complex populations, where these cell types represent a small fraction of the cell mixture. Historically, the information known about a cell lineage is correlated with its abundance and thus these rare cell types often contain new information for uncharacterized cell types. Whereas scRNA-seq methods can identify these rare cell populations, they provide only a glimpse of the RNA expression patterns in rare cells because of the detection bias for highly expressed RNAs. Moreover, because the mRNAs from these rare cells represent a small fraction of the total library, increasing the sequencing depth is not an efficient way to learn more about these cells. More complete analysis of their expression might identify e.g., cell surface markers that could be used to isolate these rare cell populations.

Recently an approach termed DART-seq was developed that enables acquisition of both global and targeted gene expression information in a single experiment (Saikia et al. 2018). In DART-seq, gene-specific probes are ligated to oligo-dT terminated Drop-seq beads (Macosko et al. 2015), enabling both oligo-dT-primed and site-specific cDNA synthesis during reverse transcription. This approach is valuable if the mRNAs of interest are known *a priori* to provide increased coverage for specific mRNAs.

Here we developed “transcriptome resampling” to address limitations of single-cell RNA sequencing. Many single-cell RNA sequencing platforms have been developed (**Table S1)** and all of them incorporate a unique DNA sequence to mRNAs derived from a single cell. We reasoned that this sequence could serve as a molecular handle to isolate RNAs derived from a cell of interest, and that these isolated RNAs could be resequenced to higher depth to interrogate the transcription profile of targeted cells. Moreover, this same principle could be applied to isolate RNAs by their unique sequences, enabling their detection in a second round of DNA sequencing. By physically isolating RNA-derived cDNA fragments, we find that transcriptome resampling can more deeply interrogate RNAs in specific cells, or can be used to determine whether specific mRNAs are expressed across cells in a mixture.

## RESULTS

### A strategy to recover individual transcriptomes from scRNA-seq libraries

The cDNA molecules in single-cell RNA sequencing libraries contain unique oligonucleotide barcodes that are used to associate a sequencing read to its cell of origin (**Fig. 1A,B**). Because these barcodes are typically long (11-19 continuous or discontinuous base pairs, depending on the platform, **Table S1**) and are present in every molecule in the library, we reasoned that these sequences could be used as sites for oligonucleotide hybridization, enabling selective recovery of specific molecules from a pool of molecules containing many different barcodes.

**FIGURE 1.**
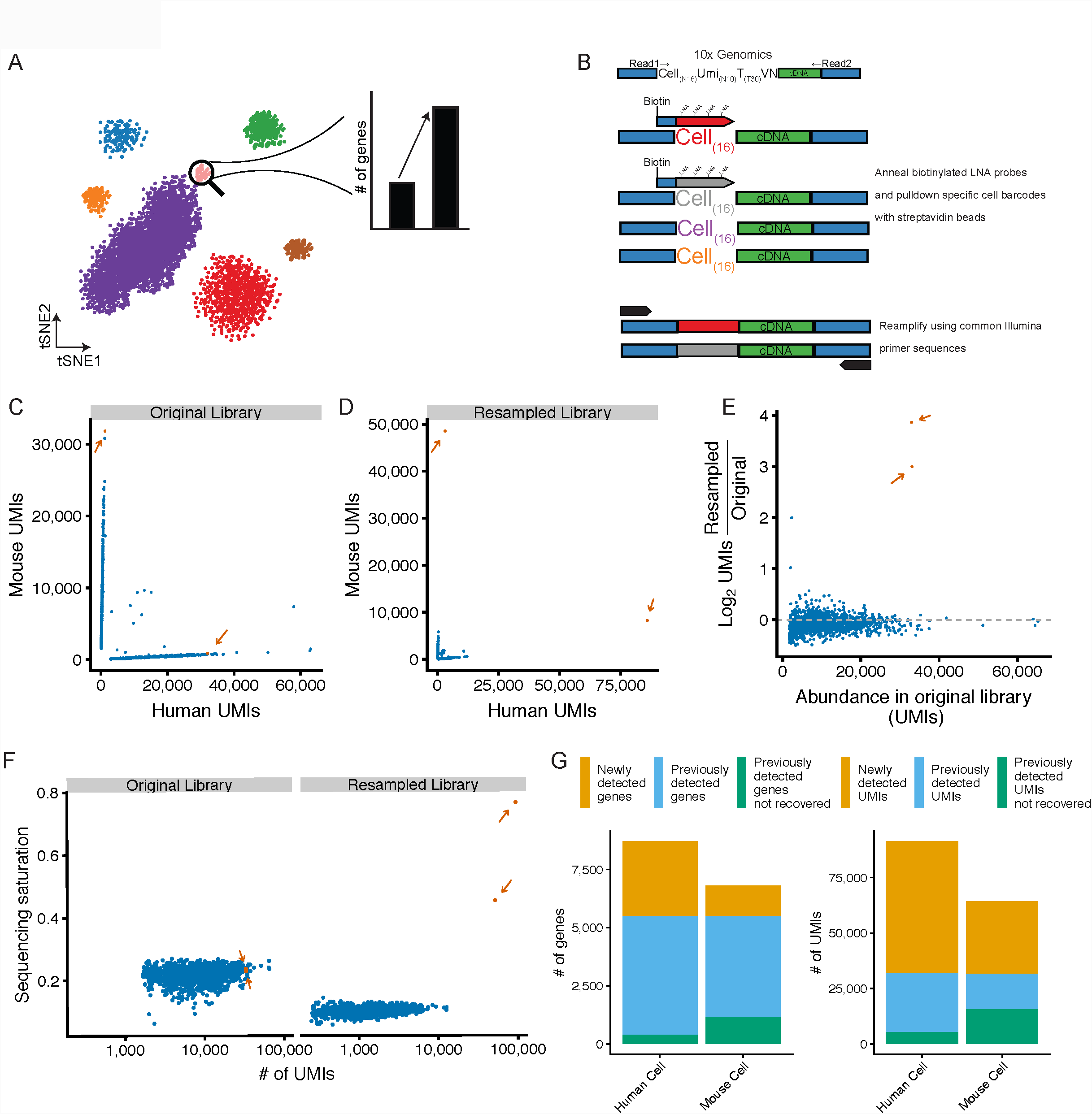
Resampling specific cell transcriptomes from pooled single-cell RNA-seq libraries. **A.** Resampling single cell libraries from rare cell populations to enable deeper characterization of a targeted cell type. **B.** Schematic of Locked Nucleic Acid (LNA) hybridization-based approach to enrich mRNAs from targeted cells from 10X Genomics single-cell mRNA-seq libraries. **C.** Species specificity of cells recovered from a 10X Genomics 3′ end gene expression scRNA-seq library containing a 1:1 mix of mouse (NIH-3T3) and human (293T) cells. Orange dots and arrows (*n* = 2) indicate cells selected for resampling; blue dots (*n* = 1,505) are untargeted cells. **D.** Species specificity of cells from the resampled library. Colors are are the same as in **C. E.** Enrichment of targeted libraries after resampling. The y-axis plots the log_2_ enrichment of UMIs normalized by the size of the entire scRNA-seq library. Colors are the same as in **C. F.** Sequencing saturation, as defined by 1 minus the ratio of the number of UMIs to the number of reads, per cell for resampled and untargeted cells in the original scRNA-seq library or after resampling. Colors are the same as in **C. G.** Number of genes (left) or UMIs (right) in the resampled cells that are either newly detected by resampling (orange), previously detected in the original library (blue), or previously detected in the original library but not found after resampling (green).

As proof-of-principle, we resampled a mouse and a human cell transcriptome from a mixed-species single cell RNA-seq library. We generated a 3′ end single cell library on the 10X Genomics Chromium platform with a 1:1 mixture of mouse NIH-3T3 fibroblasts and human 293T cell lines (**Fig. S1**). After sequencing and analysis of this library, we selected a single mouse cell and a single human cell and designed oligonucleotide probes to target their unique cell barcodes (16 nt) and a short common region (4 nt) 5′ of the cell barcode for each cell library (**Fig. 1B, Table S1**). To increase the specificity of hybridization, we incorporated locked nucleic acid (LNA) nucleotides at 6 sites in the oligo, increasing the T_m_ to 74 °C. Biotin was added to the 5′ end of the oligo to enable purification of hybridized DNA using streptavidin beads. These probes were hybridized with PCR amplified library DNA. Following streptavidin purification and elution, the enriched libraries were reamplified with PCR primers containing common Illumina sequences and sequenced.

After resampling, the targeted cell barcodes were the most abundant barcodes detected in the sequencing libraries (**Fig. 1C,D**). The number of UMIs for the resampled cells increased by 1.56-fold and 2.85-fold for mouse and human, respectively, while non-targeted cells were largely depleted at the expense of the resampled sequences (**Fig. 1E**). A useful measure of complexity in single-cell mRNA sequencing libraries is “saturation”, which is an estimate of the number of single mRNAs (as measured by UMI counts) captured from each cell in the experiment. The original single cell RNA-seq libraries were sequenced to an average saturation (i.e., the proportion of UMIs observed for a given cell at a given sequencing depth) of 21.75% +/- 2.4, and after resampling the saturation for the selected cells increased to 45.44% (mouse) and 76.63% (human) (**Fig. 1F**). Finally, we examined the number of both genes and UMIs recovered in each resampled cell and found largely the same genes and UMIs previously observed in each cell (**Fig. 1G and S2A**).

The novel UMIs recovered in the resampled cells had diverse sequence content compared to previously detected UMIs from the same gene, indicating that the novel UMIs were not simply due to resampling artifacts (**Fig. S3A**). UMIs recovered in the resampled transcriptomes were largely assigned to the expected species, although the species purity decreased by 6.06% (human cell) and 2.23% (mouse cell) upon resampling, likely reflecting increased detection of free RNA molecules present in each droplet (**Fig. S4A**). In addition, increased sequencing depth correlates to decreased species purity in our original scRNA-seq library as well as in publicly available datasets (**Fig. S4B,C**). Overall these data demonstrate the feasibility of resampling individual cell transcriptomes using LNA-based hybridization and that the rate of misassignment of reads to resampled cells is low relative to the gains achieved by resampling.

### Isolation and resequencing of rare cell transcriptomes

We next sought to more fully characterize transcriptomes derived from rare cells in a complex cell population. We generated a 10X Genomics 3′ end scRNA-seq library from a sample of peripheral blood mononuclear cells (PBMCs) taken from a healthy human adult (**Fig. 2A**). Megakaryocytes are present at ∼0.1% in the bone marrow (Nakeff and Maat 1974), have been found to populate the lung (Lefrançais et al. 2017), and are rare in a typical PBMC sample from a healthy person. We found that megakaryocytes represented 2.2% of the PBMCs as judged by expression of the megakaryocyte marker *PF4* (**Fig. S5**). We selected four megakaryocytes for resampling to further characterize the transcriptomes of these cells. After hybridization and resequencing, the barcodes for selected cells were enriched over non-targeted cell barcodes (**Fig. 2B**), were driven to a higher sequencing saturation (65.21% +/- 1.1 in original, 93.86% +/- 2.8 in resampled, **Fig. 2C**), had increased numbers of cell-derived reads (typically an order of magnitude), and resulted in the detection of an additional 677.25 +/- 143.1 genes per cell (**Fig. 2D**). To assess the specificity of cell type identification after resampling, we supplemented the original scRNA-seq dataset with the resampled cell transcriptomes. The resampled transcriptomes maintained the megakaryocyte identity as based on tSNE visualization (**Fig. 2E-G**), and the nearest neighbor for each resampled cell in PCA space was the original cell transcriptome selected for resampling (**Fig. S6**). From these data we conclude that transcriptome resampling is an efficient method to efficiently isolate and more deeply analyze the transcriptome of targeted cell types.

**FIGURE 2.**
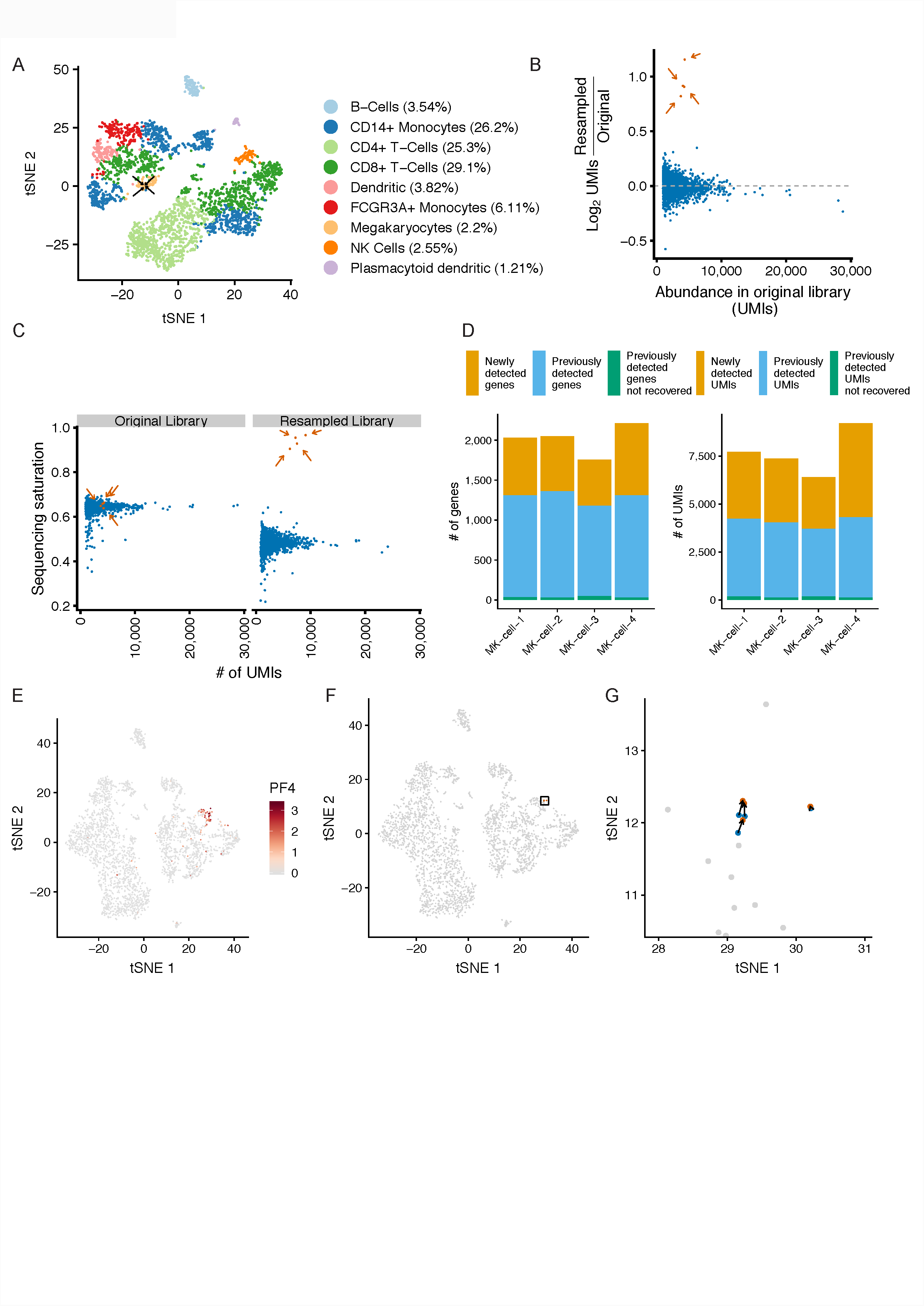
Resampling rare megakaryocytes from peripheral blood mononuclear cells. **A.** tSNE projection of a sample of peripheral blood mononuclear cells (PBMCs; *n* = 3,194 cells) with cells labels by their inferred cell type. Arrows indicate megakaryocytes selected for resampling (n = 4 resampled megakaryocytes; n = 69 total megakaryocytes). **B.** Enrichment of UMIs in targeted cells following resampling. Y-axis indicates log_2_ enrichment of UMIs normalized by library size of the entire resampled scRNA-seq library. Orange dots and arrows (*n* = 4) indicate cells selected for resampling; blue dots (*n* = 3,190) are untargeted cells. **C.** Sequencing saturation of the targeted cells in the original library and after resampling. Colors are the same as in **B**. **D.** Number of genes or UMIs in the resampled megakaryocyte (MK) cells that are either newly detected in the resampled library (orange), previously detected in the original library (blue), or previously detected in the original library but not found after resampling (green). **E.** tSNE projection of the original scRNA-seq dataset supplemented with the resampled cells. Cells are colored by the expression (natural log) of the megakaryocyte marker *PF4.* **F.** tSNE with the original cell transcriptomes (blue), resampled transcriptomes (orange), and non-targeted cells (grey). The rectangle indicates the location of cells highlighted in panel **G**. **G.** tSNE projection of resampled cells in the region highlighted in panel **F**.

### Identification of novel marker genes from resampling data

One goal of scRNA-seq experiments is to define novel marker genes that characterize a cell population. To address the utility of transcriptome resampling for this application, we examined the gene expression patterns of the resampled megakaryocyte transcriptomes. Genes most strongly enriched in the resampled libraries were more lowly expressed in the original libraries, and also had reduced levels of sequencing saturation (**Fig. 3A**). In addition to detecting more genes, each resampled cell also contained more marker genes of megakaryocytes defined from the megakaryocyte cluster in the original library (**Fig. 3B**).

**FIGURE 3.**
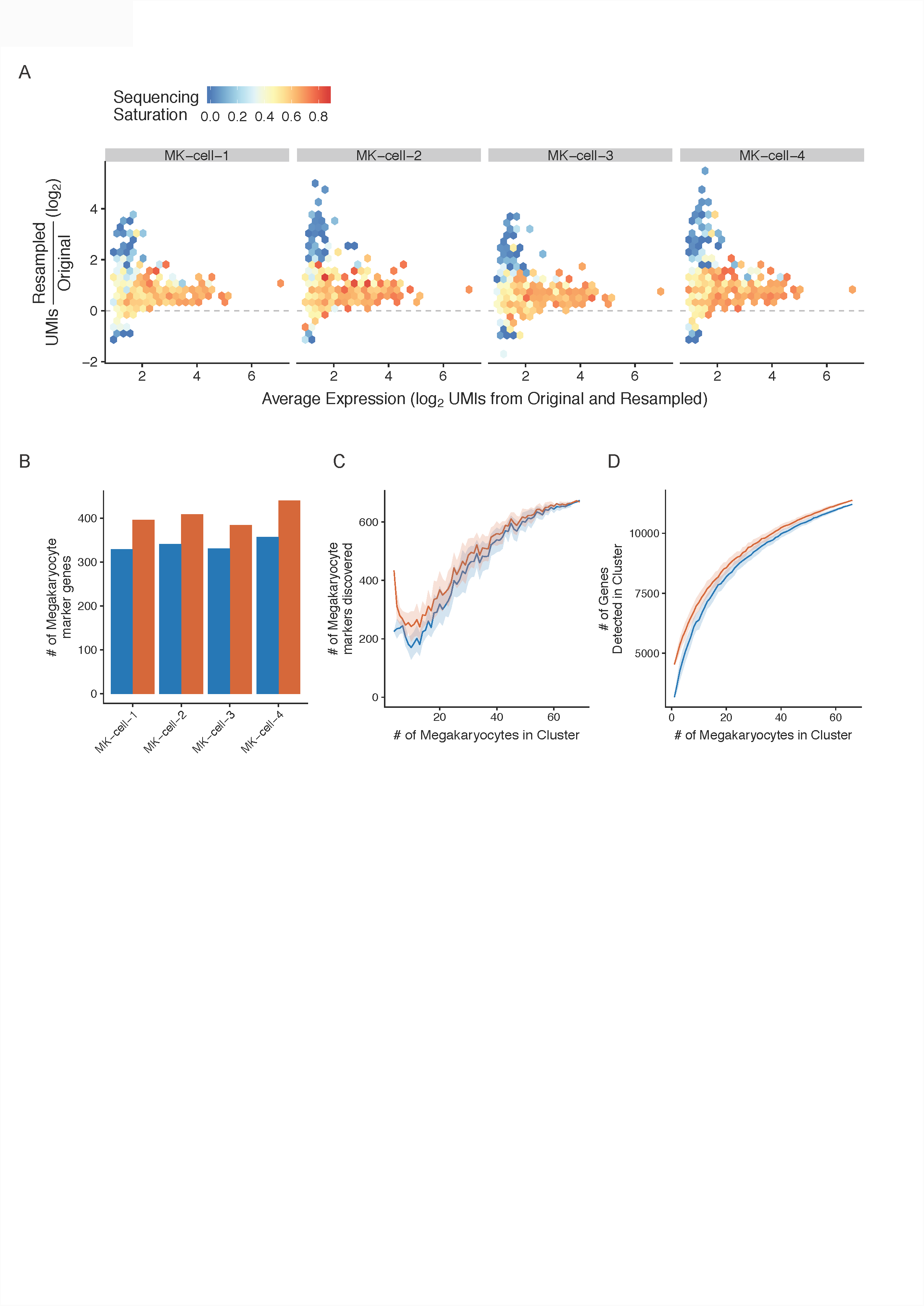
Enrichment of megakaryocyte markers in resampled cells. **A.** Relationship between sequencing saturation and enrichment of genes in the resampled libraries. X-axis indicates the normalized expression as the average of the expression values in the original and resampled libraries. Y-axis indicates UMI enrichment. Genes are binned into hexagons colored by their sequencing saturation (1 - UMIs / reads) calculated from the original library values (scale bar from 0 to 1.0). For genes not detected in the original library, the average sequencing saturation from all megakaryocytes is displayed. **B.** The number of genes recovered in each resampled cell that were defined as megakaryocyte markers in the original scRNA-seq dataset. Orange dots are for the resampled libraries; blue is from the original. **C.** Markers for megakaryocytes were calculated using the resampled cells (first set of dots), using either the UMI counts from the original library (blue dots) or the UMI counts from the resampled library (orange dots). The same procedure was repeated with increasing numbers of randomly selected non-resampled megakaryocytes supplementing the resampled cells. The random selection was repeated 10 times and the average is shown with the standard deviation displayed in the shaded areas. **D.** Number of unique genes detected in a hypothetical megakaryocyte cluster containing the cells selected for resampling from the resampled library (orange line) or the original library (blue line) with increasing numbers of randomly selected non-targeted megakaryocytes supplementing the resampled cells. The random selection was repeated 100 times and the average is shown with the standard deviation displayed in the shaded areas.

A key parameter to motivate a resampling experiment is to determine the appropriate number of cells to resample to gain additional insight into the gene expression profile of a particular cell population. In our megakaryocyte resampling experiment, we resampled 4 of the 69 (∼5%) megakaryocytes from the dataset. However, with only ∼5% of the megakaryocytes resampled, we did not find an increase in the number of genes that are differentially expressed (i.e., marker genes) between the megakaryocyte cluster and other clusters in the dataset (674 marker genes in the original library; 669 in the resampled). The resampled cells have lower normalized gene expression values (**Fig. S7**) due to increased gene diversity, particularly for highly expressed genes, resulting in small decreases in the number of novel markers detected in the resampled data. We next investigated the relationship between the number of resampled cells in a cell population and the ability to detect new cell-specific markers. We computed differentially expressed genes between the four megakaryocyte cells selected for resampling and all other non-megakaryocyte cells in the PBMC dataset, and then iteratively supplemented the four cells with additional megakaryocyte cells until all of the megakaryocytes were present in the cluster (**Fig. 3C**). At small cluster sizes, the resampled cells provided additional power to detect novel markers, but the relative increase in markers detected decreased with increasing cluster size.

Finally, we examined the relationship between the number of resampled cells and the overall number of genes recovered in the megakaryocyte cluster after resampling. Supplementing the cluster with the resampled libraries increases the overall number of genes recovered; however, at higher cells numbers the relative increase in genes detected is smaller (**Fig. 3D**). Overall, we increased the total number of detectable genes in the megakaryocyte cluster from 11,209 to 11,377. These results demonstrate that the resampled libraries allow for increased recovery of marker genes in a cell population, but the contribution of any specific resampled cell will be influenced by the initial size of the cell population sampled and the increase in the number of recovered genes in resampled cells.

### Specificity of cell barcode targeting

We considered whether hybridization-based isolation of cell-specific transcriptomes might lead to stochastic enrichment of other, non-targeted cells, so we examined the level of cross-reactivity between hybridization probes and non-targeted cell barcodes. For both the human / mouse cell and megakaryocyte experiments, we compared the level of enrichment for targeted and non-targeted barcodes with their propensity for cross-hybridization using a Smith-Waterman alignment-based score and their Hamming distance (**Fig. S8**). For the targeted human and mouse cells, we found that the cognate cell barcode was the highest alignment score and lowest Hamming distance to the hybridization probe (**Fig. S8A**). We also found two non-targeted cell barcodes with elevated enrichments above two-fold and we currently cannot explain why they are enriched. It is possible that this apparent enrichment is due to stochasticity in the low numbers of UMIs recovered for these cells. Alternatively, it is possible that these barcodes may have been enriched due to hybridization to intervening sequence in the cDNA, which is invisible in the experiment because we collected short reads from either end of the amplicon (**Fig. S8A)**. We performed a similar analysis for targeted megakaryocytes and found that for all four cell barcodes targeted by hybridization, the cognate barcodes had the highest alignment scores and lowest Hamming distances of any recovered barcodes (**Fig. S8B**), and we did not find any enrichments for non-targeted barcodes.

### Enhanced recovery of TCR mRNAs from single cells

Another common scRNA-seq application is profiling VDJ-rearranged B and T-cell receptor sequences. We applied LNA-based hybridization method to resample individual cells to recover additional VDJ rearranged receptor sequences. A TCR receptor enrichment library was prepared from the 5′ end Jurkat cell library using targeted PCR with primers specific for the TCRA and TCRB mRNAs (**Fig. 4A**). We selected two cells for which the *TCRB* chain was assembled, but in which the *TCRA* chain had not been successfully assembled. After resampling we detected enrichment for the targeted cells (**Fig. 4B**), and were able to assemble full-length *TCRA* chain in these cells after resampling (**Fig. 4C and Fig. S9**). The resampled cells had additional read coverage that spanned the consensus Jurkat *TCRA* and *TCRB* chains (**Fig. 4D**). These results demonstrate that resampling can be applied to recover single cell VDJ-rearranged TCR sequences from libraries that did not yield fully assembled TCR receptor genes.

**FIGURE 4.**
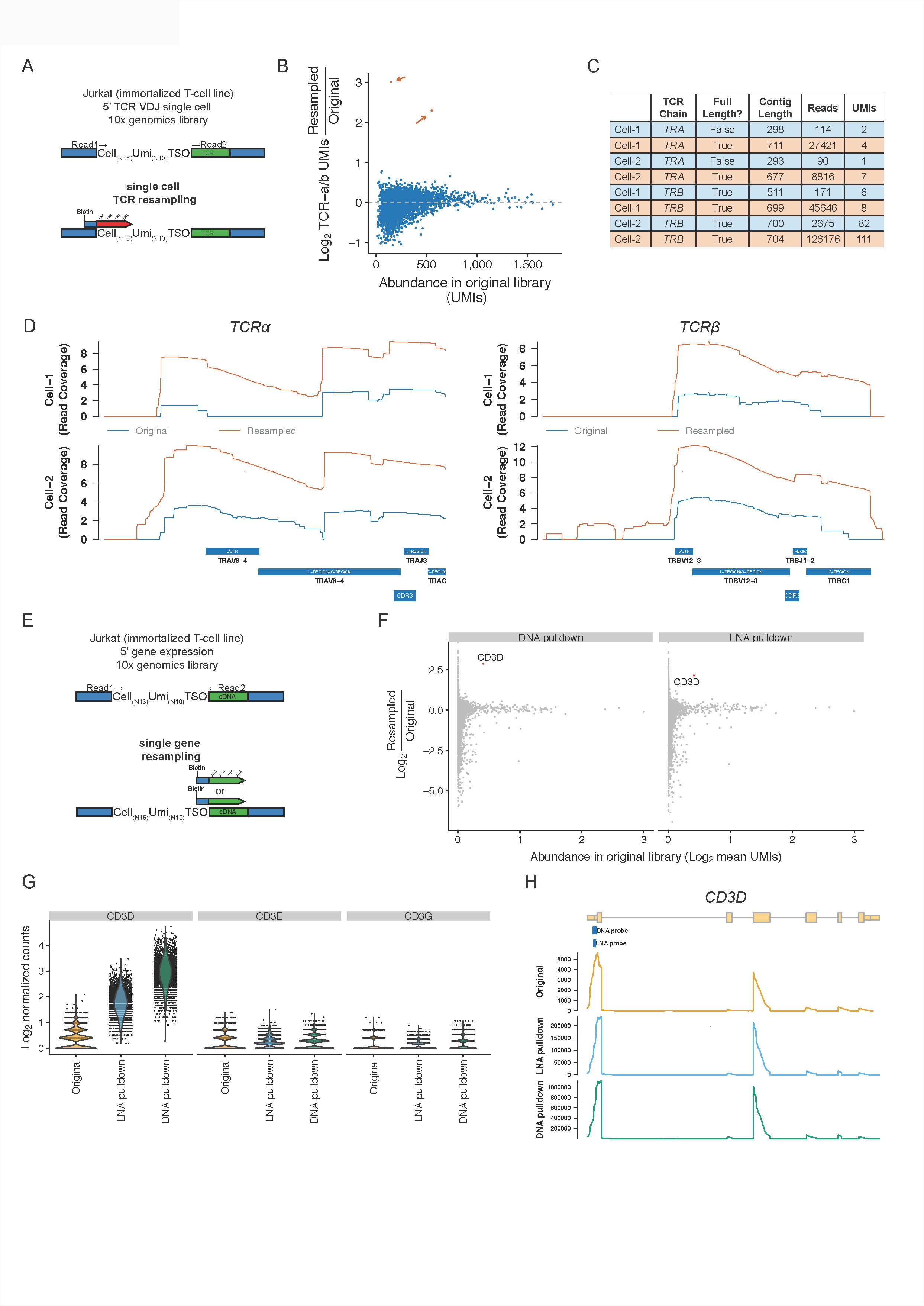
Resampling individual mRNAs and TCR sequences. **A.** Schematic of hybridization method applied to resampling single cell VDJ sequences from a 5’ end VDJ enrichment library. **B**. Enrichment of UMIs derived from the *TCR alpha* or *TCR beta* chains for the targeted cells (resampled cells; *n* = 2, shown in orange). **C.** Summary statistics of the TCR assemblies from the original (blue) or resampled libraries (orange). *Note that UMIs with less than 10 reads were excluded from the assembly and are not included in the displayed UMI and read counts. **D**. Read coverage (Log_2_ scale) across the consensus Jurkat *TCR-alpha* and *TCR-beta* sequences in the original library (blue) or the resampled library (orange). **E.** Schematic of hybridization method applied to resampling a single mRNA from a 5’ end gene expression library. Hybridization probes were designed with LNA nucleotides or with only DNA. **F.** Average fold-change of normalized UMI counts across all cells after resampling with a DNA only oligo or an LNA oligo targeting *CD3D*. X-axis indicates the normalized expression as average of the expression values in the original and resampled libraries. **G.** *CD3* expression per cell for each *CD3* chain. **H**. Read coverage across *CD3D* in the original or resampled libraries.

### Isolation and resequencing of targeted mRNA subsets

The hybridization-based resampling method can in principle be applied to isolate arbitrary oligonucleotide sequences in a single cell RNA-seq library. A common challenge with single cell RNA-seq libraries is the low cellular detection rate of low or moderately expressed genes due to stochastic capture of mRNA molecules from cells. These “gene dropout” events (i.e., false negative mRNA identifications) can prevent the identification of cells expressing key marker genes or trangenes designed to identify cell populations. We therefore examined the suitability of the resampling method to enrich specific mRNA sequences and enable interrogation of the expression of specific mRNAs across all cells.

As a proof of principle, we generated a non-saturated 5′ end gene expression library on the 10x genomics platform from Jurkat cells (an immortalized human T cell line). An initial round of sequencing showed that the mRNA for delta chain of the CD3 T-Cell coreceptor (*CD3D*) was only detectable in 59.7% of cells despite being highly expressed in this cell line (Barretina et al. 2012). To increase the cellular detection rate of the *CD3D* mRNA, we designed hybridization probes that target the 5′ end of *CD3D* mRNA (**Fig. 4**). We considered several points in the design of probes to the *CD3D* mRNA. When designing probes against an mRNA sequence there are fewer sequence constraints than with cell barcodes with fixed position and length, and therefore longer DNA-only probes might be used to increase the probe Tm. Both LNA and DNA hybridization probes directed to the 5′ ends of *CD3D* mRNA were designed and used to isolate *CD3D* fragments from 5′ gene expression libraries. We first identified a region with high read coverage in the original library near the annotated *CD3D* transcription start site. We then used a oligonucleotide probe design tool (Primer3) to pick candidate ∼25 nt sequences for LNA probes; we also used these sequences a basis for DNA hybridization probes, but included another 10-15 bases on each side of the LNA probe candidate sequences. Next, we used BLAST to rule out probes with poor E-scores or long stretches of off-target complementarity in human DNA. Finally, we selected probes with GC content >50% (especially at their 3′ ends), T_m_ values between 75 and 85 °C, and a low propensity form secondary structures at the predicted T_m_. In addition to an LNA containing probe (20 nt, T_m_ = 71 °C), we also generated a DNA only probe (40 nt, T_m_ = 66 °C) to determine whether LNAs would be necessary for specific hybridization (**Fig. 4E**).

After resampling with the LNA and DNA probes we observed specific enrichment of the *CD3D* mRNA (**Fig. 4F-G**), increasing the cellular detection rate of *CD3D* from 59.7% to 100% of cells. In addition, other CD3-associated mRNAs (*CD3E* and *CDEG*) were not enriched after resampling (**Fig. 4G**). After resampling, the read coverage profiles across the *CD3D* mRNA remained similar, indicating minimal bias in the read coverage introduced by resampling (**Fig. 4H**). These results demonstrate the utility of using LNA and DNA hybridization probe for resampling specific mRNA species.

### Recovery of individual transcriptomes by targeted PCR

We additionally tested a PCR-based strategy to recover targeted single cell transcriptomes. The 10x genomics platform uses cellular barcodes that are Hamming distance 2 apart, which cannot be reliably distinguished by standard PCR approaches. In contrast, other platforms such as the Wafergen iCell8 system, have more diverse barcodes (Hamming distance of 3), with limited numbers of barcodes detected per experiment (1,000s of detectable barcodes versus 100,000s of barcodes in 10x genomics or DropSeq libraries). The Wafergen iCell8 libraries contain an 11 nucleotide cell barcode (**Fig. S10A**).

To investigate the utility of a PCR-based approach, we designed PCR primers to anneal to cell barcodes and recover the transcriptomes of single cells from a scRNA-seq library. We tested three PCR approaches to recover single cell transcriptomes. The first strategy used standard DNA primers, the second strategy incorporated a 5′ biotin to enable stringent purification of PCR products, and the final strategy additionally incorporated phosphorothioate linkages into the terminal three 3′ nucleotides to prevent 3′-to-5′ exonucleolytic cleavage by the proofreading Phusion DNA polymerase (See **Table S2** for primer design).

To test these PCR strategies, we selected either 10 (standard and biotinylated approach) or 5 cells (phosphorothioate approach) that spanned ∼100-fold in sequencing coverage and were derived from either human cells (from a breast cancer tumor xenograft) or mouse cells (host mouse bone marrow derived cells). We performed two rounds of PCR with low cycle numbers to enrich the libraries. We performed 14 cycles of amplification in individual reactions, then either pooled the resulting PCR products for the standard approach, or for the biotinylated and phosphorothioate approach purified the PCR products using AMPure purification followed by streptavidin magnetic beads to remove unamplified library material. Lastly we performed a second round of PCR to incorporate library indexes and sequences required for flow cell clustering. We observed 7.7, 8.6, and 19.7-fold enrichment for the targeted barcodes in the resampled sequencing libraries, for the standard, biotinylated, and biotinylated with phosphorothioate approach respectively (**Fig. S10C-E**). Both the standard PCR and biotinylated purification approach were enriched for many non-targeted sequences. In contrast, barcodes were specifically enriched over non-targeted barcodes by the third PCR approach. These results indicate that incorporation of phosphorothioates is necessary to achieve specific amplification, and suggests that 3′-to-5′ exonucleolytic activity of the Phusion polymerase is responsible for the observed non-specific amplification.

A caveat of using a direct PCR resampling approach is the potential for rebarcoding of similar barcode sequences. Non-specific hybridization to similar barcode sequences would result in the amplification of non-targeted cell barcode sequences. These off-target sequences would be misclassified to a resampled cell due to the PCR primer sequence rebarcoding the original off-target cell barcode. To investigate the extent of rebarcoding in these libraries we assessed the proportion of UMIs that were found in multiple cells in the original library and the resampled library. In the LNA-based hybridization method, the fraction of UMIs that were detected in multiple cells in the original library did not appreciably increase in the resampled libraries (0.76% +/- 0.26 shared in original library, 1.80% +/- 0.47 shared in resampled libraries) (**Fig. S3B**), indicating that the novel UMIs recovered after resampling were not likely derived from other cells. In contrast, resampling with the PCR method resulted in increases in the proportion of UMIs that were found in multiple cells (0.39% +/- 0.26 to 3.52% +/- 1.75 for the phosphorothioate approach) (**Fig. S10G**). Additionally the percentage of UMIs assigned to the correct species was lowered after resampling with the PCR approach (for the phosphorothioate approach, 91.26% +/- 8.8 in the original library compared to 76.66% +/- 27.5 for the resampled library) (**Fig. S10H**). These results demonstrate that a direct PCR based resampling approach can result in undesired off-target rebarcoding, which for the purposes of examining single cell gene expression profiles makes the PCR approach less desirable that the LNA-based approach.

## DISCUSSION

Here we demonstrate how resampling of individual transcriptomes from scRNA-seq libraries provides richer information for selected cell and mRNA populations. One consideration in the design of probes for the resampling approach is the design and structure of barcode information in single-cell mRNA sequencing libraries. We leveraged the contiguity of the cell barcode in 10X Genomics and Wafergen libraries to design probes and primers that can effectively target molecules by hybridization. Such an approach could also be used for other platforms where a contiguous cell barcode is synthesized on each bead (e.g., Drop-seq, **Table S1**) (Macosko et al. 2015). The 10X Genomics Chromium platform uses a small set of fixed cell barcodes (737,280 in version “737K-august-2016.txt”) and each of these have on average ∼13 sequences within Hamming distance of 2. For 10X Genomics libraries, we opted for the hybridization approach as it is less likely to lead to “recoding” of cells in the library. Moreover, because cell barcodes are short in 10X Genomics libraries (16 nt), we used locked nucleic acid probes that target only the barcode region. We found that resampling by PCR is not ideal because primers designed against one barcode might misprime on and amplify a related barcode, causing inclusion of mRNA from an unrelated cell in the resampled transcriptome.

Before embarking on a resampling experiment one must ensure that a single-cell mRNA sequencing library is sufficiently complex as to yield additional information after targeted resampling and resequencing. Libraries with high overall saturation observed in the first round of sequencing are unlikely to benefit from resampling because most UMIs have already been observed. We selected libraries with less than ∼66% saturation to maximize the information gained from specific cells. New methods that increase cell numbers in single-cell experiments (Stoeckius et al. 2017; Gehring et al. 2018) will benefit from transcriptome resampling because as cell numbers increase, DNA sequencing becomes limiting and fewer reads are recovered per-cell from these more complex libraries. Transcriptome resampling may enable an initial low-depth examination of many cells followed by more targeted analysis of defined populations. As an example, we increased sequencing coverage for four megakaryocytes (out of of library with 3,194 detected cells; **Fig. 2A**) between six and 20-fold after resampling. The current recommended maximum for captured cells in the 10X Genomics workflow is 10,000 cells; therefore if a cell were resampled from this larger population, we would expect and 18 to 60-fold increase in its sequencing coverage, assuming the same sequencing depth and efficiency of hybridization for cell-specific probes.

Both LNA and DNA probes performed well for *CD3D* mRNA, increasing the percentage of cells with detected expression from 59.7% to 100%. We also found a DNA probe that targeted the same region of *CD3D* provided enhanced recovery of UMIs associated with *CD3D* mRNAs (**Fig. 3C**). DNA probes are less expensive than LNA probes (∼$60 per probe for biotinylated DNA versus ∼$200 per LNA probe) and may be more cost effective when targeting multiple mRNAs. We envision pooling subsets of mRNA-specific probes to more fully characterize gene expression programs in cells from specific contexts (e.g., interferon-stimulated (Schneider et al. 2014) or stress-response (de Nadal et al. 2011) expression programs).

Our expectation was that upon resampling we would observe a uniform increase in the number of UMIs across genes expressed at different levels. Instead, we found that resampling tends to recover UMIs from genes expressed at low levels (**Fig. 3A**), indicating that UMIs from highly expressed genes may already approach saturation and that resampling is useful to characterize the expression of genes expressed at low levels. Moreover, inspection of UMI recovery for cells before and after sampling showed that, while lowly expressed genes are recovered a higher rates, the UMIs for highly expressed genes are proportionally decreased, possibly reflecting the true abundance of the molecules in the library that is only observed after resampling (**Fig. S7**).

In principle, transcriptome resampling might be used to query other features of mRNA expression and processing. For example, full-length mRNAs might be studied in greater detail by isolating molecules from libraries containing full length cDNA (e.g., Smart-Seq2 (Picelli et al. 2013)). This approach may be valuable for single-molecule sequencing approaches, where read numbers are limiting. However, probe-based mRNA isolation is subject to some caveats. Several biological processes create variation at mRNA 3′ ends including alternative polyadenylation and 3′ UTR splicing. In addition, 3′ end libraries have a large degree of internal mispriming at genomically-encoded poly(A) streches (Shepard et al. 2011), potentially rendering a large proportion of the cDNA from a given mRNA unable to be captured using a 3′ end targeted probe. It is possible that 5′ gene expression libraries have a more homogeneous representation of a given mRNA 5′ end, enabling the design of hybridization probes that target a majority of mRNA isoforms.

We anticipate that the recovery of individual transcriptomes will facilitate characterization of rare cell populations identified in scRNA-seq experiments. However, the structure of DNA barcodes in a scRNA-seq library impacts the generality of the resampling approach. We applied resampling to libraries wherein cell barcodes are encoded by a contiguous region of DNA. As such, a single hybridization probe can specifically recover information for an individual cell. Other library designs employ discontinuous cell bar codes (e.g., sci-RNA-seq (Cao et al. 2017)); here the information needed to associate an mRNA with a single cell is present at different sites in the molecule (i.e., a linker sequence in addition to the two library indices). In this case, enrichment for a portion of the cell barcode would like provide additional information for the cell of interest, but would also enrich for other unrelated cells because some cell barcode information is distant to the site of hybridization.

Resampling could also be used to recover molecules from other types of complex single cell libraries. Single-cell ATAC-seq and DNA-seq have been used to probe chromatin accessibility and copy number variation in individual cells (Cai et al. 2015; Cusanovich et al. 2015; Buenrostro et al. 2015). Because the amplicons in these libraries have structure similar to scRNA-seq libraries with a cell barcode and UMI, one could resample cells with interesting chromatin properties from these mixed populations. Detection of regulatory regions and transcription factor footprints is highly dependent on read coverage (Landt et al. 2012) and deeper sequencing of recovered libraries could provide more insight into gene regulation than can be gained from the mixed population, possibly enabling interrogation of how promoters and enhancers are correlated in accessibility in single-cell ATAC experiments or providing increased depth of coverage for targeted domains in single-cell Hi-C (Ramani et al. 2017).

## METHODS

### Cell isolation and single cell RNA-seq generation

For the mouse / human experiment, NIH:3T3 and 293T cells were grown in standard media and mixed at a 1:1 ratio prior to capture. For the mouse xenograft experiment, an estrogen receptor positive patient derived xenograft primary cell line (UCD65 (Kabos et al. 2012)) was labeled with luc-eGFP using lentivirus (Hanna et al. 2016). Cells were xenografted into NOD/SCID/IL2rg^-/-^ mice via intracardiac injection to generate disseminated metastases or injected into the mammary pad to generate a primary tumor. Cells were isolated from the primary cell line, a primary fat pad tumor, a brain metastasis and a bone metastases. Contaminating murine bone marrow cells copurified with human tumor cells isolated from the bone metastasis. Single cells were then captured using the iCell8 system from WaferGen Bio-Systems.

For the Jurkat libraries, Jurkat cells were thawed from a frozen vial, grown for one passage in RPMI-1640 + 10% FBS media and diluted to 500 cells / µL for capture on the 10X Genomics Chromium controller. For the megakaryocyte experiment, blood from healthy controls between ages 15 and 55 was collected in heparinized tubes and peripheral blood mononuclear cells (PBMC) were isolated by density gradient centrifugation using Ficoll to obtain the buffy coat.

Wafergen libraries were prepared according to the manufacturers instructions. 5′ and 3′ gene expression libraries were prepared on the 10X Genomics Chromium controller and libraries prepared according to manufacturers protocol. TCR enrichment libraries were generated using the 10X Genomics 5′ library construction and VDJ enrichment kit.

### PCR primer design and synthesis

For 10X Genomics cell barcode pulldown libraries, 5′ biotinylated oligos with 4 base pairs of consensus sequence followed by the 16 basepair barcode with LNA bases added every 3rd base. These 6 LNA basepairs allow for increased binding specificity of the barcode sequence. For 10X Genomics gene pulldown libraries, 5′ biotinylated oligo probes were designed against selected CD3D using primer3 (Untergasser et al. 2012). For LNA probes, LNA bases were added every 4rd base. Regions to target were selected based on inspection of read coverage of the 5′ UTR of the *CD3D* gene (See **Fig. 4H** for genome browser tracks). LNA molecules were purchased from either Exiqon or Qiagen. For Wafergen libraries, 5′ biotinylated oligos with 17 basepairs of consensus sequence followed by the 11 basepair barcode were designed containing 3 phosphorothioate bonds on the three terminal 3′ positions to prevent degradation. All oligonucleotides used in this study are described in **Table S2**.

### Library preparation and sequencing

10X Genomics Illumina libraries at 2nM concentration were diluted 20 fold and PCR amplified with Truseq Pcr F and R oligo for 14 cycles with 1 unit of Phusion. PCR amplification was confirmed by gel electrophoresis and the remaining sample was Ampure XP purified at a ratio of 1.8x bead/PCR volume. Samples were eluted in 10µl and molarity was determined using a Qubit. 10 µl of a 100 nM library was mixed with 10 pmol LNA molecule in annealing buffer (10 mM Tris 8.0, 50 mM NaCl) and heated to 98° for 10 minutes followed by slow cooling to 59°-64° depending on the LNA and held overnight at the annealing temperature. Annealed DNA was added in equal volume to washed Dynabead M270 streptavidin beads at the annealing temperature. Beads were incubated for 15 minutes with gentle shaking. Beads were washed five times in binding buffer at the annealing temperature. Hybridized molecules were eluted in 20 µl of elution solution buffer (50 mM NaCl, 0.1 mM EDTA) by heating for 10 minutes at 100° and centrifuging briefly to pellet the streptavidin-bound LNA molecules (which can inhibit PCR). The elution was performed two times to ensure all the DNA was removed from the LNA. 10µl of streptavidin purified input DNA was added to PCR using primers Truseq PCR F and R for 8 cycles. PCR products were ampure purified, analyzed by Tapestation D1000, quantified by the Qubit, and sequenced on an Illumina MiSeq or Nova Seq 6000.

Wafergen Illumina libraries at 2 nM concentration were diluted 20 fold and PCR amplified with 1 unit of Phusion and the cell specific biotinylated primer and Nextera read 2 sequencing primer to specifically amplify the sequences from the selected cells for 14 cycles. PCR amplification was confirmed by gel electrophoresis and the remaining sample was purified with Ampure XP beads at a ratio of 1.8x bead/PCR volume. Samples were eluted and added in equal volume to prepared Dynabead M270 streptavidin beads. Beads were incubated for 15 minutes at room temperature with gentle rotation. Beads were washed 5x in bind and wash buffer and eluted in 20 µl of water. 10 µl of streptavidin-purified input DNA was added to PCR using PEPCRPrimer 1.0 and Nextera N703 for 9 cycles. PCR products were AMPure purified, confirmed on the Agilent TapeStation D1000, quantified by Qubit and submitted for sequencing.

### Data analysis

We provide our data analysis pipeline and custom scripts in a github repository (https://github.com/rnabioco/scrna-subsets). Single cell RNA-Seq libraries were preprocessed to append the read 1 sequence to the paired read 2 read id followed by quality trimming and poly(A) tail removal from read 2 using cutadapt (v. 1.8.3) (Martin 2011). Reads were next aligned with STAR (Dobin et al. 2013) to either the human genome assembly (hg38) for the PBMC experiments or a genome with both human (hg38) and mouse (mm38) sequences for all other experiments. Sequence headers in the human/mouse combined genome were prefixed with either an “H_” or “M_” to designate human or mouse references respectively.

Following alignment, BAM files were processed to extract the cell barcode and UMI sequences into tags (CN and BX) within the BAM file. The cell barcode was error corrected against a list of cell barcodes, either as known well barcodes (Wafergen experiments), or generated from the original 10x genomics single cell libraries processed with the 10x genomics software cellranger (v. 2.1.1). Cell barcodes within an edit distance of 1 of the known barcodes were considered valid cell barcodes and corrected to the known barcode. Alignments not designated as multi-mapping that overlapped distinct exonic features were tagged in the BAM file (subread v. 1.6.0) (Liao et al. 2014). Gencode v25 annotations was used for human data, and a union reference containing Gencode v25 and mouse v11 was used for human/mouse datasets. UMIs per gene were enumerated per cell using umi-tools (v 0.5.3) (Smith et al. 2017) using the directional method to disambiguate similar UMI sequences.

tSNE projections were generated using the Seurat R package (Butler et al. 2018). Briefly, PCA analysis was performed on scaled, log-transformed, library-size-normalized UMI matrices using variable gene sets. PCA was used to reduced the dimensionality and tSNE projections were generated with a perplexity of 30. Graph-based clustering was performed to identify clusters using the first 18 principal components. Markers per cluster were identified using a Wilcoxon rank sum test. K-nearest neighbors were identified using the top 20 PCA dimensions using the RANN R package.

### Availability of data and material

DNA sequencing data are available from NCBI GEO under accession GSE119428. A reproducible software pipeline (including Snakemake (Köster and Rahmann 2012) pipeline and R Markdown documents) is available at https://github.com/rnabioco/scrna-subsets. Processed data is available from zenodo at https://zenodo.org/record/1405579.

## ACKNOWLEDGMENTS

We thank Srinivas Ramachandran for comments on the manuscript, and Katrina Diener and Todd Woessner for expert technical assistance. This work was supported by the RNA Bioscience Initiative at the University of Colorado School of Medicine and the National Institutes of Health (R35 GM119550 to J.R.H.). Single-cell RNA sequencing libraries and Illumina sequencing was performed in the University of Colorado Cancer Center Genomics Shared Resource (P30 CA046934).

## SUPPLEMENTAL FIGURE LEGENDS

**Figure S1.**
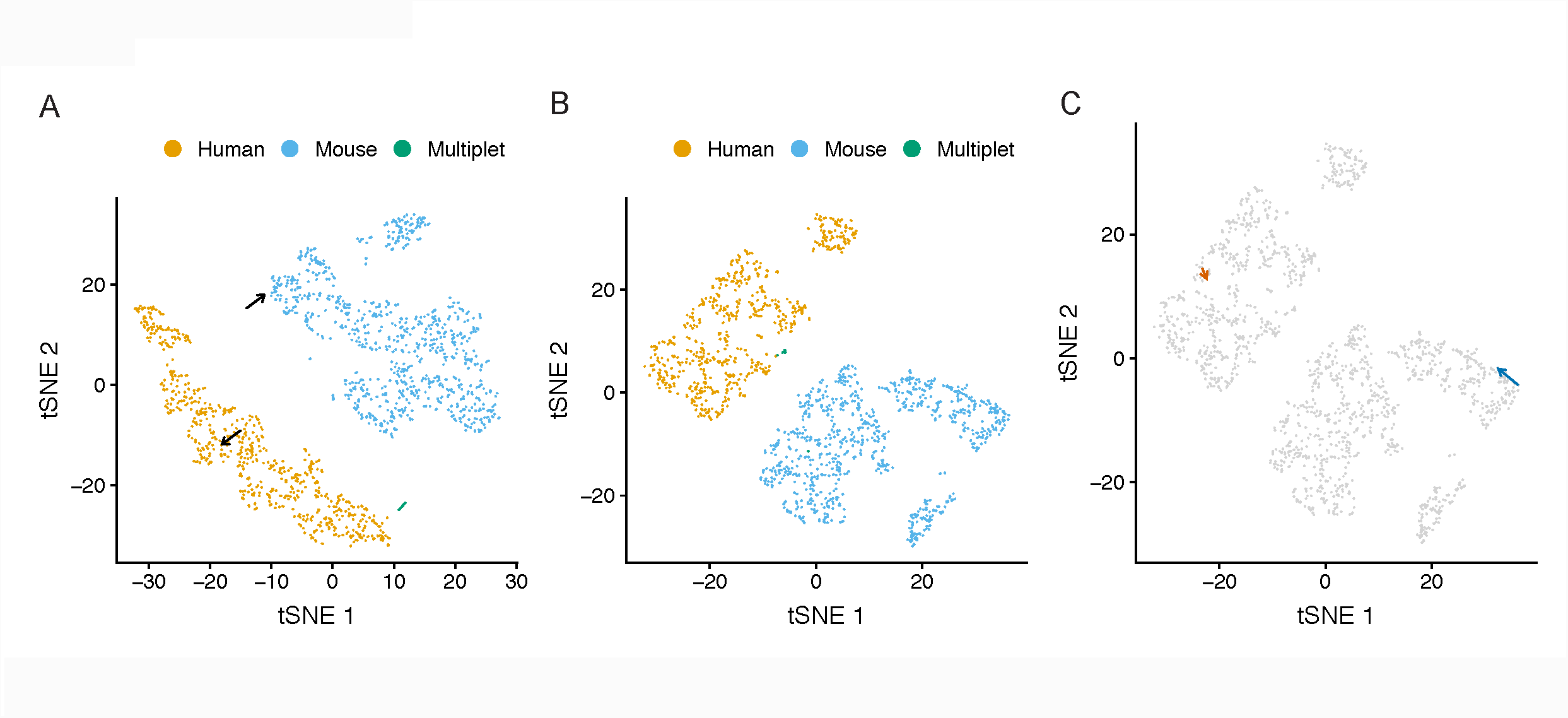
Resampling single mouse and human cells from a NIH3T3:293T scRNA-seq library. (A) tSNE projection of a scRNA-seq library prepared from a 1:1 mix of NIH3T3 mouse and 293T human cells. Arrows indicated cells selected for resampling. (B) tSNE projection of the NIH3T3:293T library supplemented with the resampled cells (C) Same tSNE projection as B with arrows drawn from the original cell to the resampled cell.

**Figure S2.**
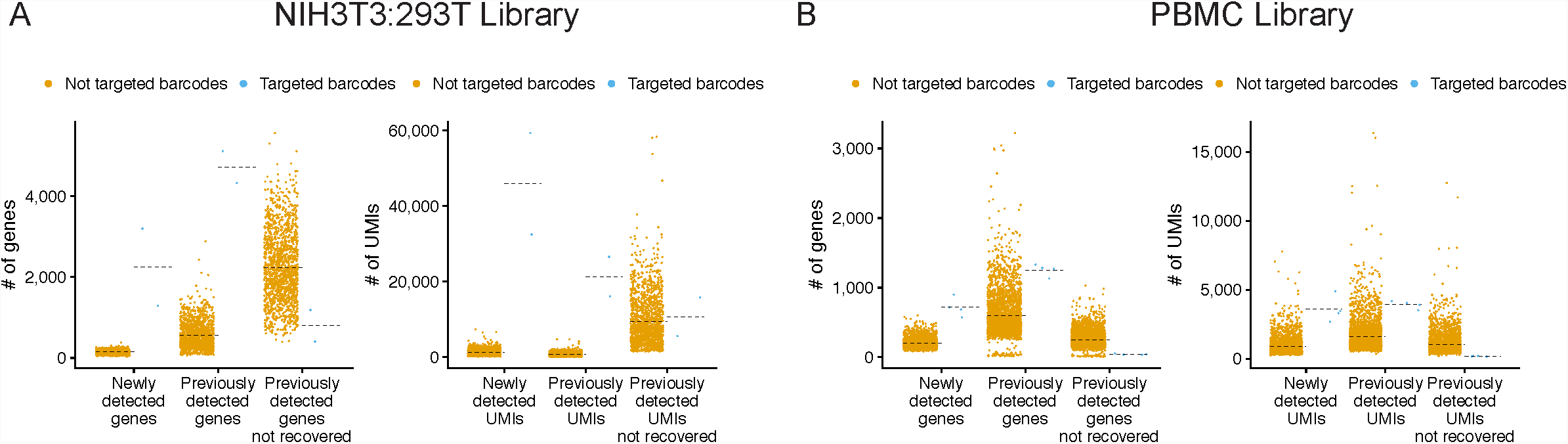
Resampled libraries have a higher proportion of new UMIs compared non-targeted resampled libraries. Number of genes and UMIs recovered that are newly detected, were previously detected, or were previously detected but not recovered in the resampling library. The targeted barcodes are shown in blue and the non-target barcodes are shown in orange. Mean values are indicated with a dashed line. (A) Mouse human mix library or (B) PBMC library.

**Figure S3.**
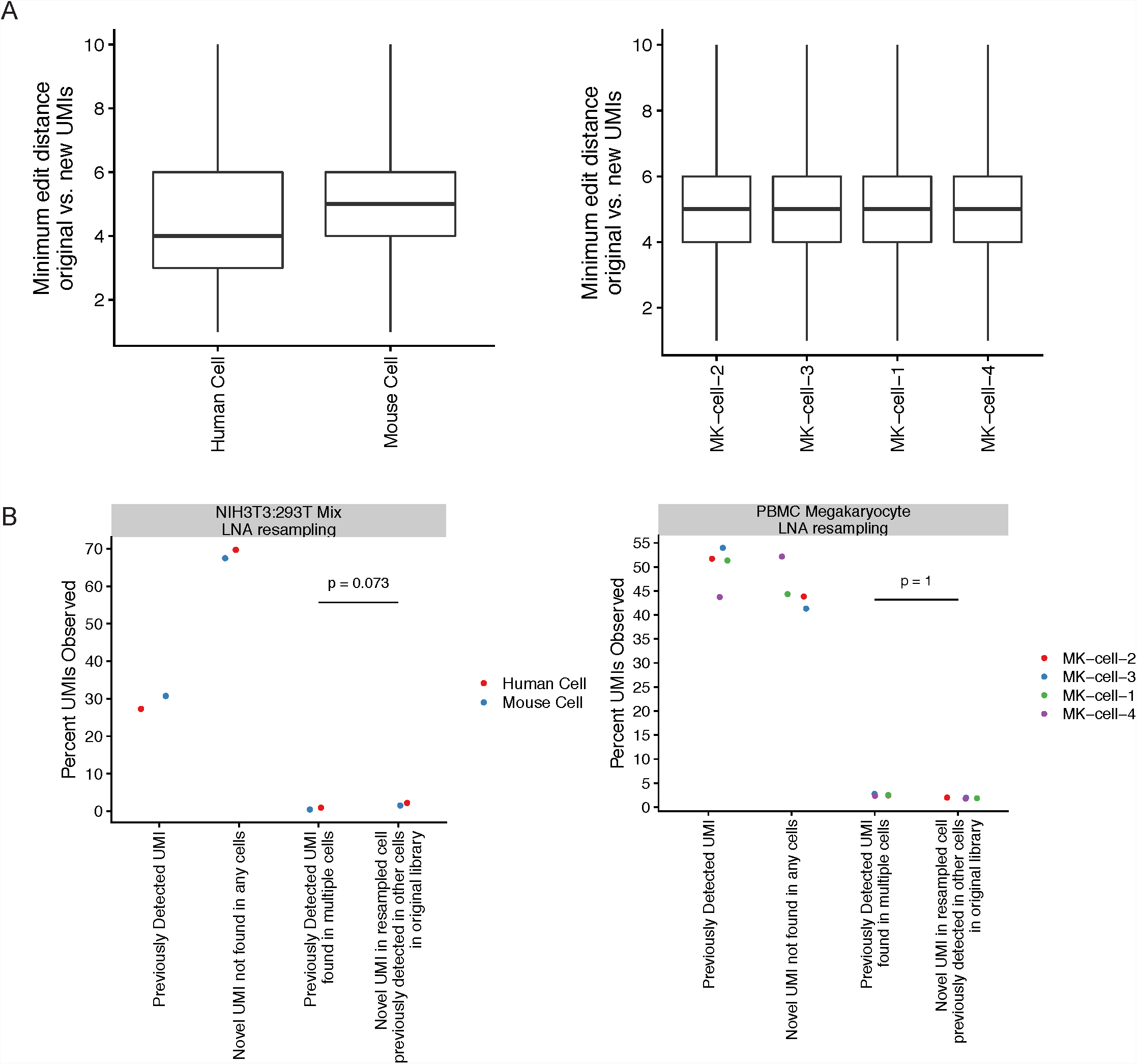
UMIs recovered by LNA hybridization are not likely derived from other non-targeted cell libraries. (A) Minimum hamming distances for novel UMIs recovered in the resampled libraries compared to all original UMI sequences. (B) Analysis of the percent of UMIs in the resampled NIH3T3/293T mix that either had been previously detected in the original library of the targeted cell, are novel UMIs not seen in any cells in the original data, had previously been observed in the original library in multiple cells, are novel UMIs in the resampled cells that had previously been observed in non-targeted cells in the original library. P-value derived from one-way student t-test testing for increased percentage of novel UMIs in the resampled cells that had previously been observed in non-targeted cells.

**Figure S4.**
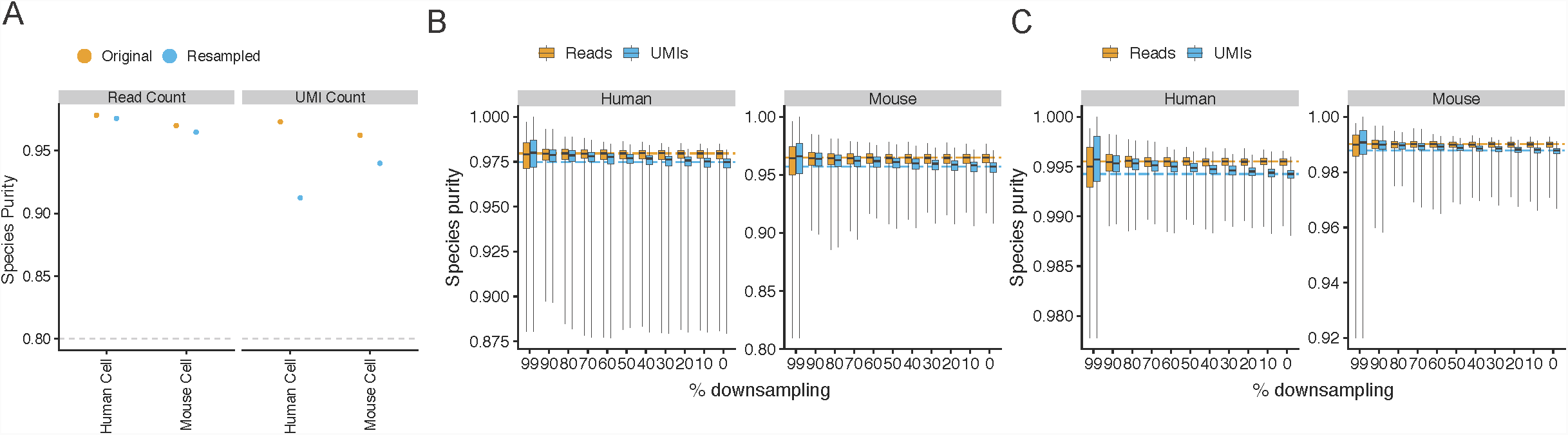
Reduced sequencing depth correlates with higher species-specificity in mixed species samples. (A) Species purity of the resampled mouse and human cells in the original or resampled libraries. Dashed lined indicates cut-off for assigning a cell as either mouse or human (B) Mapped reads were downsampled to the indicated percentage of original mapped reads. UMIs and reads remaining after downsampling were enumerated and the species purity was calculated as the ratio of # of reads/UMIs in the assigned species for the cell to the total number of human and mouse reads/UMIs. (B) NIH-3T3:293T cell sample described in this study (C) NIH-3T3:293T cell sample dataset produced by 10x genomics (https://support.10xgenomics.com/single-cell-gene-expression/datasets/2.1.0/hgmm_1k).

**Figure S5.**
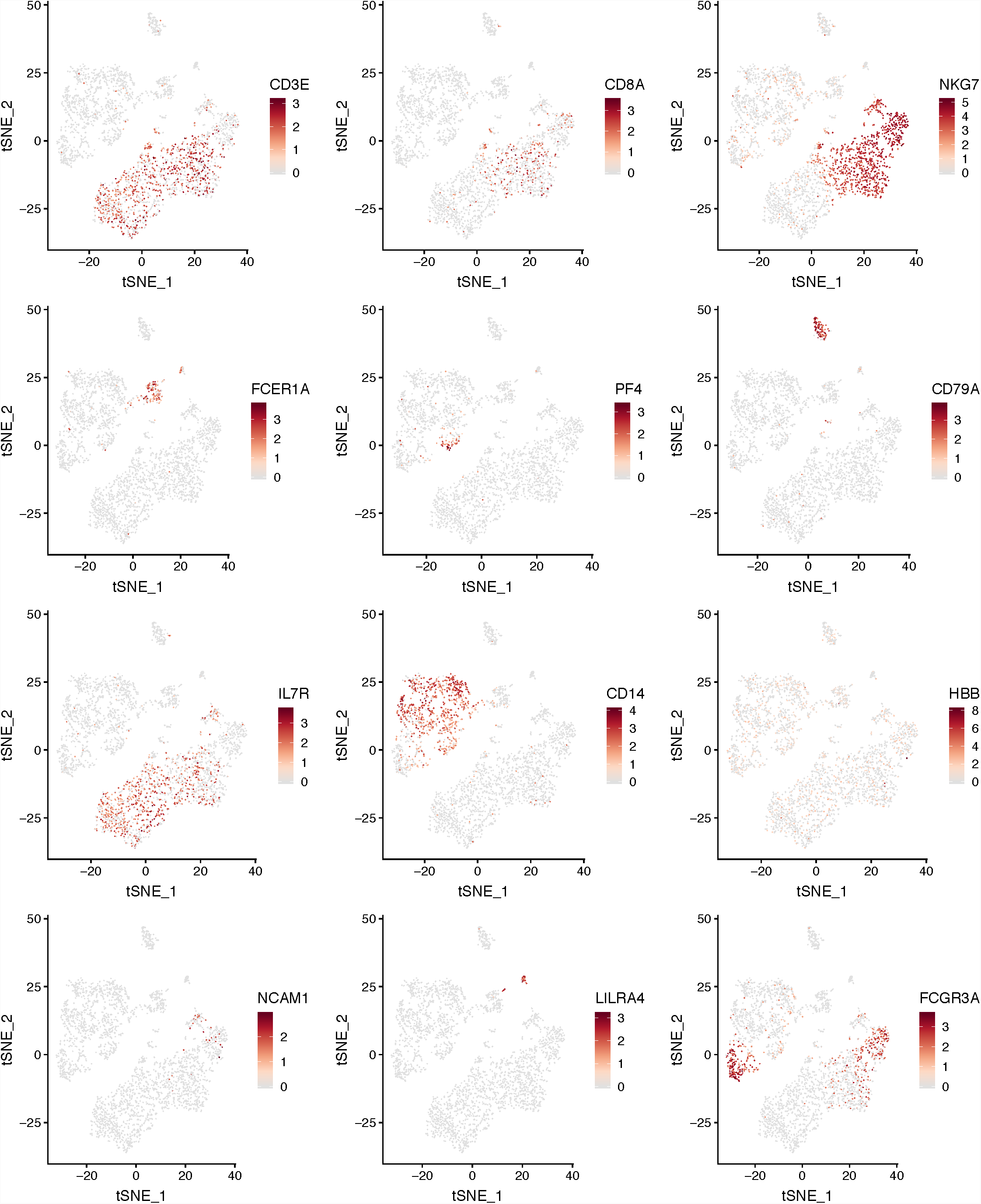
Marker genes identifying cell types present in the PBMC dataset. Expression of marker genes for cell types identified in PBMCs. Expression is shown on the natural log scale.

**Figure S6.**
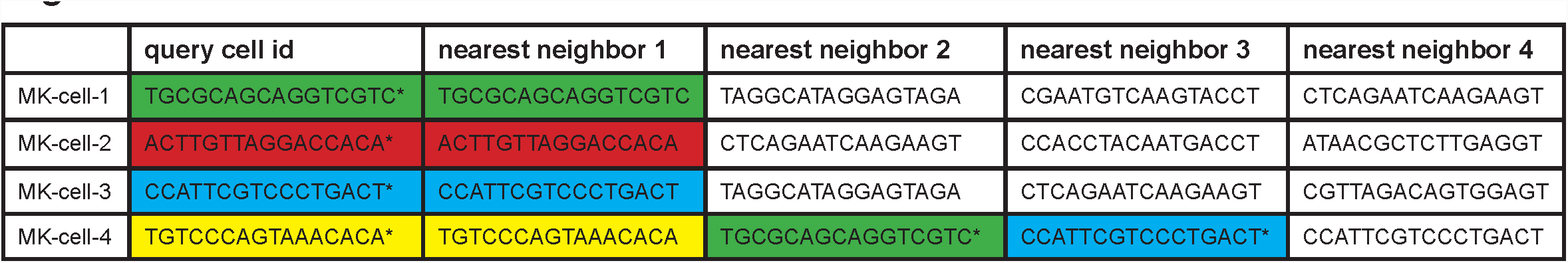
The nearest neighbors to each resampled cell in the PBMC dataset are the original not resampled cells. Asterisk indicates a resampled cell library. The original library was supplemented with the resampled libraries to evaluate the nearest neighbors of the resampled libraries. Nearest neighbors were computed in the first 20 PCA dimensions. All top nearest neighbors recovered were also classified as Megakaryocytes.

**Figure S7.**
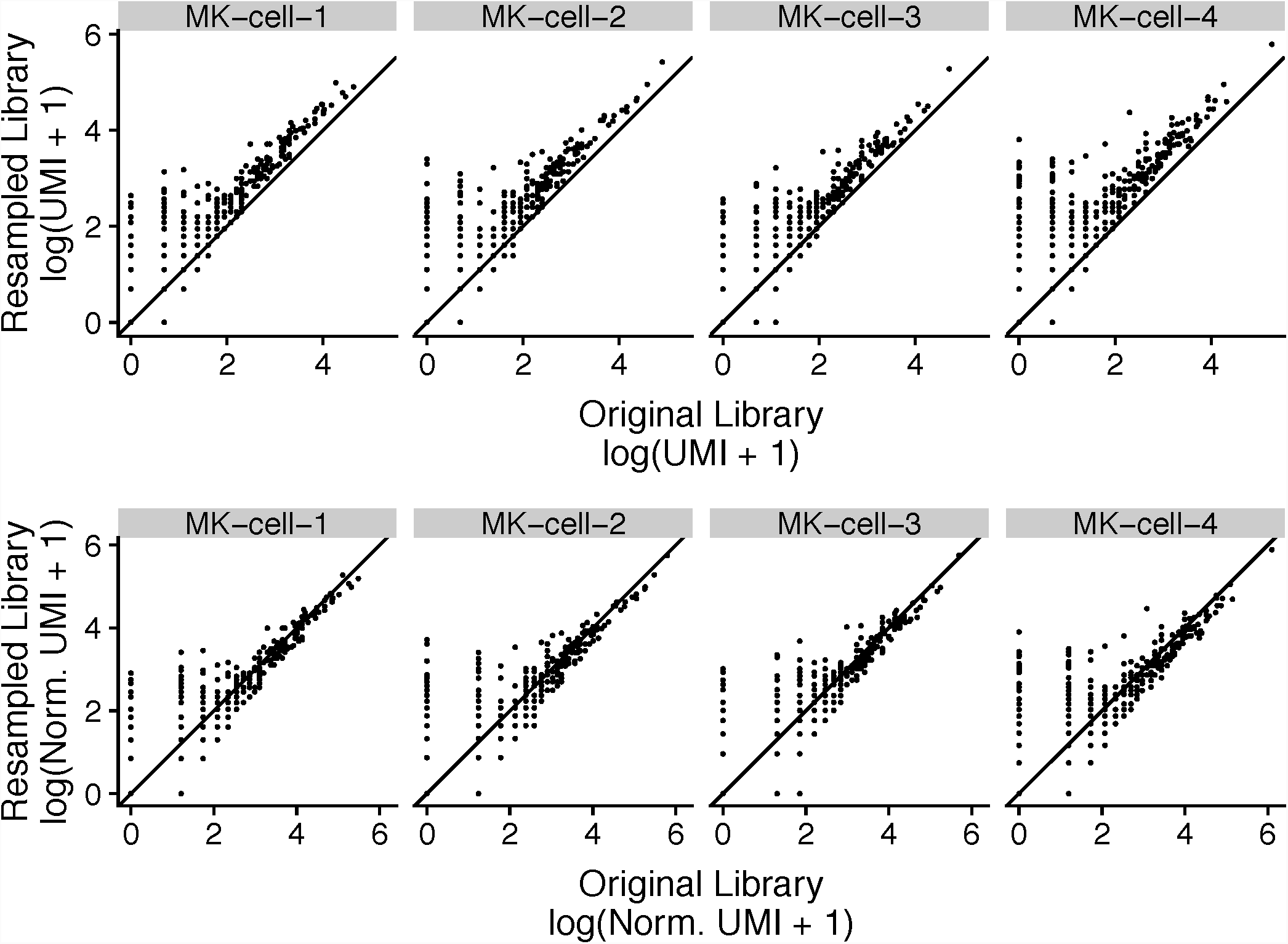
Library size normalization reduces resampled library gene expression values. UMI count distributions for cells selected for resampling. Top panel shows unnormalized UMI counts. The bottom panel show library size normalized UMI counts when the resampled cells are included within the original scRNA-seq UMI count matrix.

**Figure S8.**
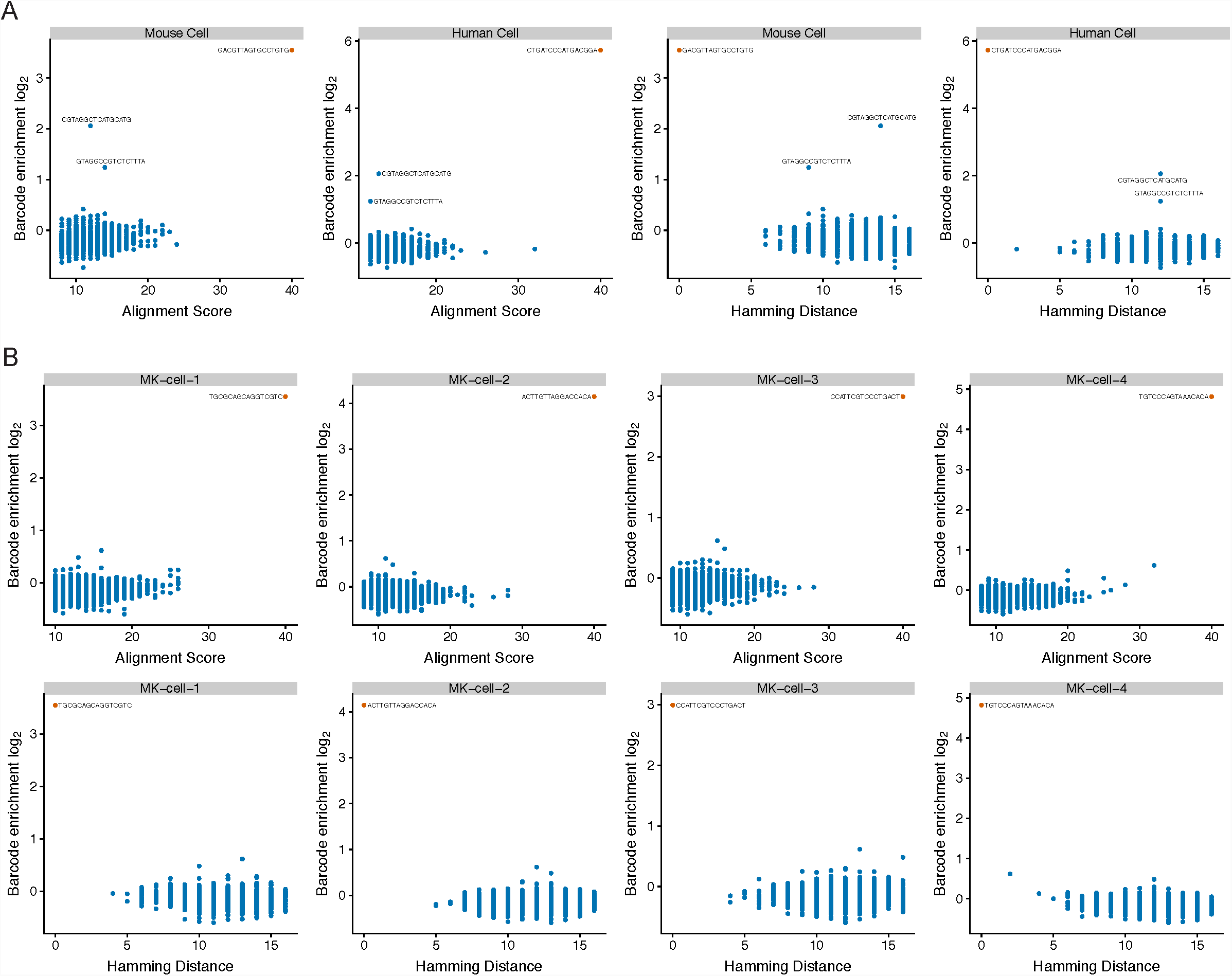
Barcode enrichment by LNA hybridization is specific. Alignment scores and hamming distances computed against each resampled cell barcode are shown plotted against the barcode enrichment (library sized normalized read counts in log scale). Alignments were performed using the full LNA probe sequence (cell barcode + 4nt of 5’ illumina sequence), against all cellular barcodes prefixed with 16 nuceotides of the known illumina sequences 5’ of the cell barcode. Alignment scores were computed using Smith-Waterman local alignment and a score of 40 is a perfect alignment. Hamming distances were computed against the cell barcode region of the LNA probe and all cellular cell barcodes. Any barcodes sequences enriched over 2 fold are indicated. Cell barcodes for the the NIH3T3:293T mixture experiment in (A) and PBMC experiment as shown in (B).

**Figure S9.**
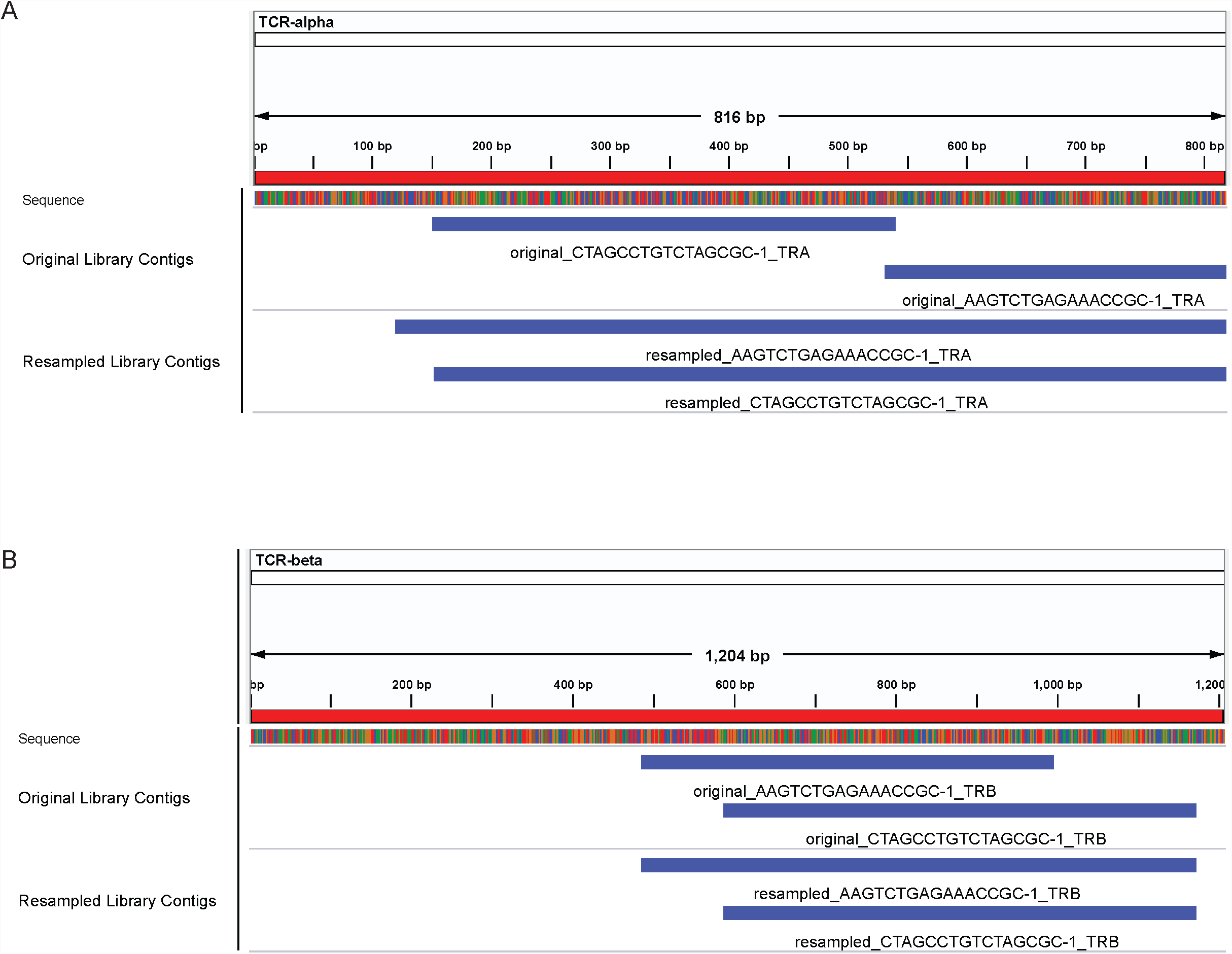
TCR alpha receptor assembly improves in resampled cells. IGV snapshots of the contigs assembled per cell from the original libraries or the resampled libraries. TCR-alpha shown in panel A, and TCR-beta shown in panel B.

**Figure S10.**
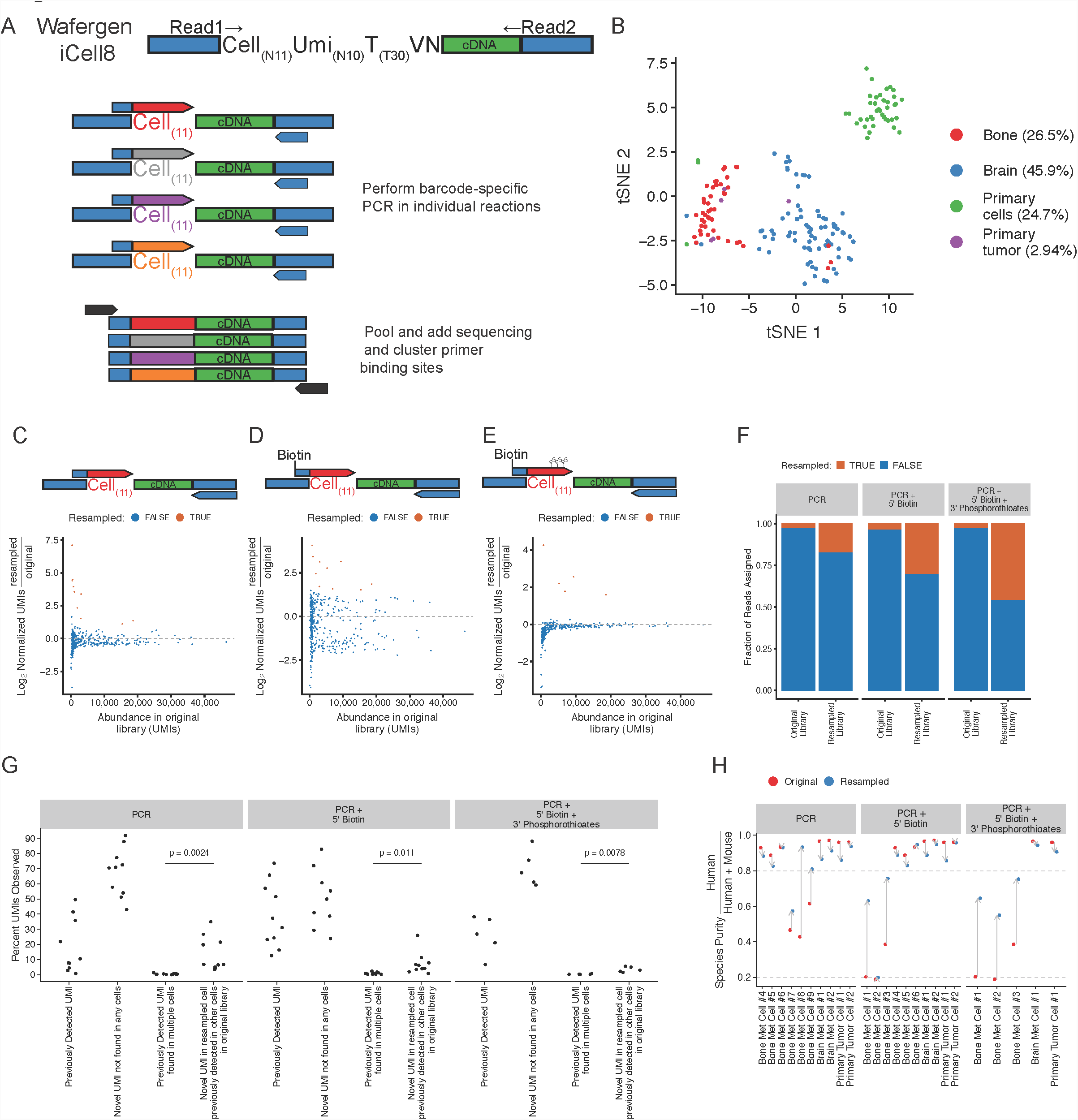
Resampling individual single cell libraries using a PCR approach. (A) Schematic of PCR only approach to enrich for single cell libraries from a Wafergen iCell8 prepared single cell library. Individual PCR reactions are performed with each cell barcode specific primer, then reaction products are pooled prior to a final PCR step to introduce illumina sequencing primer binding sites (B) tSNE projection of scRNA-seq dataset prior to resampled. A breast cancer patient derived xenograft was processed to obtain single cells from a bone and brain metastasis, a primary fat-pad derived tumor, or the original primary cell line. (C) Standard PCR based enrichment of 10 targeted cells. (D) Biotinylated primer based PCR enrichment of 10 targeted cells. Primers supplemented with biotin of the 5’ end of the primer. PCR products were immunoprecipitated with magnetic beads containing streptavidin prior to pooling in the final PCR. (E) Biotinylated primer based PCR enrichment supplemented with phosphorothioate (PS) linkages in the terminal 3’ nucleotides to prevent exonucleolytic cleavage by DNA Polymerase. 5 cells were selected for resampling. (F) Proportion of the resampled library consumed by the resampled cell libraries or other non-targeted libraries. (G) Analysis of the percent of UMIs in the resampled cells that either had been previously detected in the original library of the targeted cell, are novel UMIs not seen in any cells in the original data, had previously been observed in the original library in multiple cells, are novel UMIs in the resampled cells that had previously been observed in non-targeted cells in the original library. p-value derived from one-way student t-test testing for increased percentage of novel UMIs in the resampled cells that had previously been observed in non-targeted cells. (H) Per cell species specificity in the original libraries, or after resampling, as assessed by either enumerating the proportion of human reads per cell or human UMIs per cell. Dashed lines indicate cut-offs for classifying cells are human (80%) or mouse (20%). Arrows connect the original library values to the resampled library values.

## TABLES

**Table S1. Library designs for various single-cell RNA-Seq platforms.**

**Table S2. Oligonucleotide sequences used in this study.**

